# Exceptional yield vesicle packaged recombinant protein production from *E. coli*

**DOI:** 10.1101/2022.04.22.489142

**Authors:** Tara A. Eastwood, Karen Baker, Bree R. Streather, Nyasha Allen, Lin Wang, Stanley W. Botchway, Ian R. Brown, Jennifer R. Hiscock, Christopher Lennon, Daniel P. Mulvihill

## Abstract

We describe a novel system that exports diverse recombinant proteins in extracellular vesicles from *E. coli*. The vesicles not only compartmentalise toxic, insoluble and disulphide bond containing proteins in a soluble and functional form (e.g. DNaseI, nanobodies and IgG-fusions), but the continued release of the inducible vesicle packaged proteins into the media supports continuous isolation of protein from active culture within a micro-environment allowing stable long-term storage. This technology results in unprecedented yields of vesicle packaged functional proteins for efficient downstream processing for a wide range of applications from discovery science to applied biotechnology and medicine.

## Main Text

The ability to reprogram a cell to direct the packaging of specific molecules into discrete membrane envelopes is a major objective for synthetic biology. Controlled packaging into membrane vesicles supports the development of numerous new technologies and commercialisable products within the applied biotechnology and medical industries, including: generation of recombinant bioreactors; environmental dispersion of biomolecules; vehicles for drug delivery and vaccination, as well as providing a stable environment for isolation and storage of proteins.

We fortuitously discovered that recombinant expression of the amino terminal 38 residues of human synucleins brings about the continuous formation and release of extracellular vesicles from *E. coli* cells into the culture media (Fig. 1a). This Vesicle Nucleating peptide (VNp) interacts with the bacterial cell membrane (Supplementary Fig. 1) and induces localised curvature of the bacterial membrane that extends outwards until formation of a distinct vesicle, which is released into the growth media upon membrane scission (Fig. 1 b-d). This process occurs repeatedly without impacting upon cell viability and so drives a sustained and large-scale production of vesicles from the same cell (Fig. 1d, Supplementary Fig. 2), to support the continuous expression and isolation of recombinant proteins from media of a growth culture over a number of days, rather than only from cells harvested upon termination of the culture. Thus providing significant savings in both time and resource.

**Figure 1.**
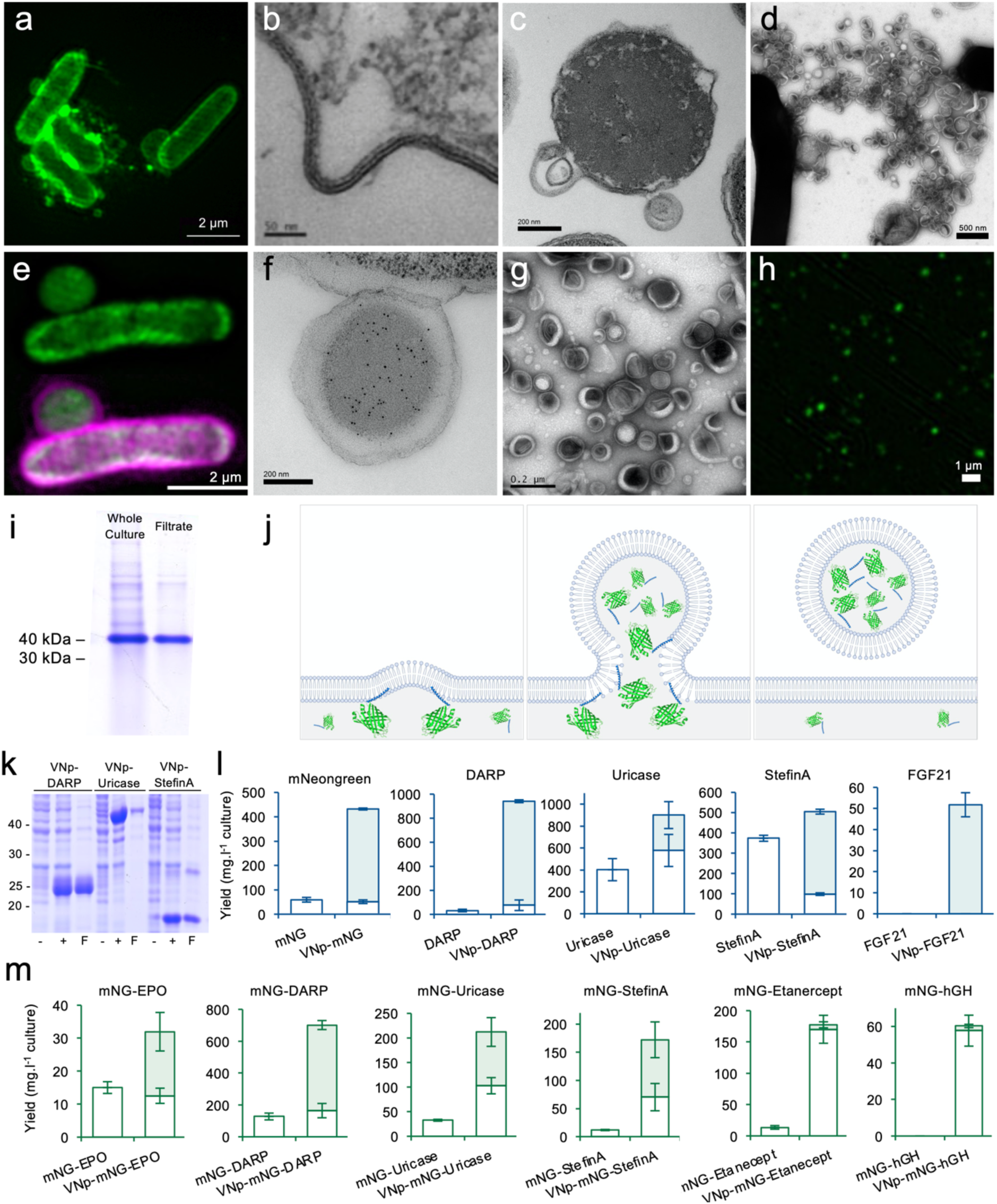
Recombinant vesicle formation. (a) OmpA-mCherry SIM fluorescence and (b & c) TEM images illustrating VNp induced membrane curvature in *E. coli*. (d) EM of vesicles generated from VNp expressing *E. coli* cells that were cultured on prepared grids. (e) mCherry (magenta) and mNeongreen (green) SIM fluorescence of VNp-mNeongreen OmpA-mCherry expressing *E. coli* cells. (f) Anti-mNeongreen immuno-EM of a section though *E. coli* associated VNp-mNeongreen induced vesicle. (g) TEM and (h) mNeongreen fluorescence images of isolated VNp-mNeongreen containing vesicles. (i) Coomassie stained gel of cell culture and filtered media of VNp-mNeongreen expressing cells. (j) Schematic of VNp induced cargo containing vesicles. (k) Coomassie stained samples of uninduced and induced cultures or filtered induced cultures of VNp-DARP, VNp-Uricase and VNp-StefinA expressing cells. (l & m) Average soluble yields per litre of culture derived from cell extracts (empty boxes) or filtered culture media (filled boxes) for each recombinant protein examined. Recombinant proteins lacked (l) or possessed (m) a fluorescent mNeongreen fusion. Errors are s.d. from ≥ 3 experimental repeats.

Fusion of sequences encoding VNp to those encoding the monomeric fluorescent protein mNeongreen^1^ led to the production and export of large VNp-mNeongreen protein vesicles into the culture media (Fig. 1 e-f). Immuno-electron microscopy confirmed the exclusive localisation of the mNeongreen cargo within the lumen of the vesicles (Fig. 1f, Supplementary Fig. 3). Low speed centrifugation and subsequent filtration with sterile 0.45 μm polyethersulfone (PES) filters efficiently and effectively isolated the vesicles from bacteria (Fig. 1g-h, Supplementary Fig. 4). Remarkably, these isolated vesicles have uniform size and provide a stable environment for effective long term protein storage of functional folded recombinant protein (Supplementary Fig. 4 & 5). The degree of purity of the fusion protein harvested by one step filtration is sufficient for a very wide range of applications (Fig. 1i), while simultaneously supporting subsequent purification after vesicle sonication, where necessary. Together these data support a model of the VNp-fusion inducing localised outward curvature of the membrane and subsequent export into the forming vesicle, which upon maturation is released into the culture media (Fig. 1j).

Thus, this system provides a simple and attractive mechanism for releasing membrane packaged recombinant proteins into the media enabling both enhanced recombinant protein production and subsequent processing. While the mNeongreen provided rapid quantification of soluble target protein exported into the media, a wider range of model biopharmaceuticals representing a range of different physical properties and expression challenges (such as membrane binding, disulphide-bond containing, or otherwise insoluble or toxic proteins) were used to test the applicability of this technology for the expression of the spectrum of molecules demanded by the life sciences community. Expression of each protein was tested as VNp, or VNp-mNeongreen amino terminal fusions and compared to the expression of equivalent non-VNp fusion proteins (Fig. 1 k-m, Table 1).

**TABLE 1:**
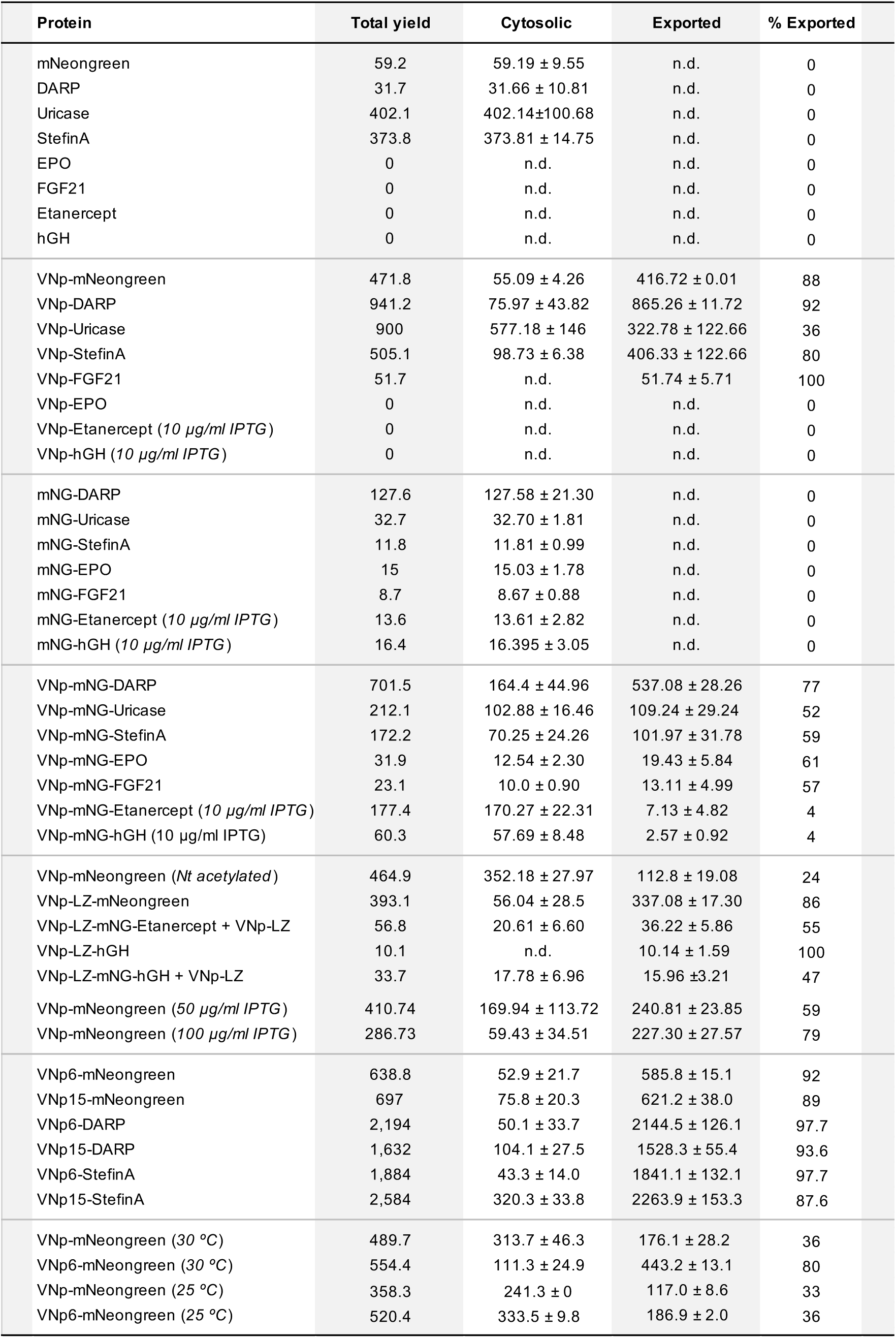
Summary of soluble protein yields. Yields measured as mg of soluble recombinant protein / litre. Cells grown in shaking flask cultures at 37 ºC with T7 promoter induced with 20 μg / ml IPTG unless stated otherwise. Average yields ± s.d. calculated from ≥ 3 independent biological repeats. n.d.: not detectable

| Protein | Total yield | Cytosolic | Exported | % Exported |
| --- | --- | --- | --- | --- |
| mNeongreen | 59.2 | 59.19 ± 9.55 | n.d. | 0 |
| DARP | 31.7 | 31.66 ± 10.81 | n.d. | 0 |
| Uricase | 402.1 | 402.14 ± 100.68 | n.d. | 0 |
| StefinA | 373.8 | 373.81 ± 14.75 | n.d. | 0 |
| EPO | 0 | n.d. | n.d. | 0 |
| FGF21 | 0 | n.d. | n.d. | 0 |
| Etanercept | 0 | n.d. | n.d. | 0 |
| hGH | 0 | n.d. | n.d. | 0 |
| VNp-mNeongreen | 471.8 | 55.09 ± 4.26 | 416.72 ± 0.01 | 88 |
| VNp-DARP | 941.2 | 75.97 ± 43.82 | 865.26 ± 11.72 | 92 |
| VNp-Uricase | 900 | 577.18 ± 146 | 322.78 ± 122.66 | 36 |
| VNp-StefinA | 505.1 | 98.73 ± 6.38 | 406.33 ± 122.66 | 80 |
| VNp-FGF21 | 51.7 | n.d. | 51.74 ± 5.71 | 100 |
| VNp-EPO | 0 | n.d. | n.d. | 0 |
| VNp-Etanercept (10 µg/ml IPTG) | 0 | n.d. | n.d. | 0 |
| VNp-hGH (10 µg/ml IPTG) | 0 | n.d. | n.d. | 0 |
| mNG-DARP | 127.6 | 127.58 ± 21.30 | n.d. | 0 |
| mNG-Uricase | 32.7 | 32.70 ± 1.81 | n.d. | 0 |
| mNG-StefinA | 11.8 | 11.81 ± 0.99 | n.d. | 0 |
| mNG-EPO | 15 | 15.03 ± 1.78 | n.d. | 0 |
| mNG-FGF21 | 8.7 | 8.67 ± 0.88 | n.d. | 0 |
| mNG-Etanercept (10 µg/ml IPTG) | 13.6 | 13.61 ± 2.82 | n.d. | 0 |
| mNG-hGH (10 µg/ml IPTG) | 16.4 | 16.395 ± 3.05 | n.d. | 0 |
| VNp-mNG-DARP | 701.5 | 164.4 ± 44.96 | 537.08 ± 28.26 | 77 |
| VNp-mNG-Uricase | 212.1 | 102.88 ± 16.46 | 109.24 ± 29.24 | 52 |
| VNp-mNG-StefinA | 172.2 | 70.25 ± 24.26 | 101.97 ± 31.78 | 59 |
| VNp-mNG-EPO | 31.9 | 12.54 ± 2.30 | 19.43 ± 5.84 | 61 |
| VNp-mNG-FGF21 | 23.1 | 10.0 ± 0.90 | 13.11 ± 4.99 | 57 |
| VNp-mNG-Etanercept (10 µg/ml IPTG) | 177.4 | 170.27 ± 22.31 | 7.13 ± 4.82 | 4 |
| VNp-mNG-hGH (10 µg/ml IPTG) | 60.3 | 57.69 ± 8.48 | 2.57 ± 0.92 | 4 |
| VNp-mNeongreen (*Nt acetylated*) | 464.9 | 352.18 ± 27.97 | 112.8 ± 19.08 | 24 |
| VNp-LZ-mNeongreen | 393.1 | 56.04 ± 28.5 | 337.08 ± 17.30 | 86 |
| VNp-LZ-mNG-Etanercept + VNp-LZ | 56.8 | 20.61 ± 6.60 | 36.22 ± 5.86 | 55 |
| VNp-LZ-hGH | 10.1 | n.d. | 10.14 ± 1.59 | 100 |
| VNp-LZ-mNG-hGH + VNp-LZ | 33.7 | 17.78 ± 6.96 | 15.96 ± 3.21 | 47 |
| VNp-mNeongreen (50 µg/ml IPTG) | 410.74 | 169.94 ± 113.72 | 240.81 ± 23.85 | 59 |
| VNp-mNeongreen (100 µg/ml IPTG) | 286.73 | 59.43 ± 34.51 | 227.30 ± 27.57 | 79 |
| VNp6-mNeongreen | 638.8 | 52.9 ± 21.7 | 585.8 ± 15.1 | 92 |
| VNp15-mNeongreen | 697 | 75.8 ± 20.3 | 621.2 ± 38.0 | 89 |
| VNp6-DARP | 2,194 | 50.1 ± 33.7 | 2144.5 ± 126.1 | 97.7 |
| VNp15-DARP | 1,632 | 104.1 ± 27.5 | 1528.3 ± 55.4 | 93.6 |
| VNp6-StefinA | 1,884 | 43.3 ± 14.0 | 1841.1 ± 132.1 | 97.7 |
| VNp15-StefinA | 2,584 | 320.3 ± 33.8 | 2263.9 ± 153.3 | 87.6 |
| VNp-mNeongreen (30 °C) | 489.7 | 313.7 ± 46.3 | 176.1 ± 28.2 | 36 |
| VNp6-mNeongreen (30 °C) | 554.4 | 111.3 ± 24.9 | 443.2 ± 13.1 | 80 |
| VNp-mNeongreen (25 °C) | 358.3 | 241.3 ± 0 | 117.0 ± 8.6 | 33 |
| VNp6-mNeongreen (25 °C) | 520.4 | 333.5 ± 9.8 | 186.9 ± 2.0 | 36 |

The VNp fusion enhanced the expression and secretion for each target protein highly effectively, and supports the expression of individual proteins ranging from less than 1 kDa (VNp-His6) to 85 kDa (VNp-mNeongreen-Etanercept) in size, as well as protein complexes as demonstrated by fluorescence from pairs of Bimolecular Fluorescence Complementation VNp-fusions^2^ within exported vesicles (Supplementary Fig. 5a). Importantly, VNp fusion enhanced the overall yield of each target protein examined, with yields of almost 1g soluble protein / litre of shaking flask culture obtained in the case for DARP (Table 1). The versatility of the system was demonstrated by the production of correctly folded (e.g. mNeongreen), membrane binding (e.g. FGF21), and enzymatically active (e.g. uricase) proteins (Supplementary Fig. 4 & 5). The vesicle isolated VNp-uricase was not only as enzymatically active as uricase purified from a cell pellet, but this activity was maintained to a higher degree by VNp-uricase stored within isolated vesicles for 2 months at 4 ºC when compared to purified protein stored at 4 ºC in buffer over the same period (Supplementary Fig. 5), highlighting the stable environment the vesicles afford their protein cargo.

The VNp fusion allows production of soluble folded proteins that are otherwise insoluble or reduce the viability of bacterial cells (e.g. DNase, Etanercept, EPO & hGH) (Table 1, Supplementary Fig. 6). In the case of the disulphide bond containing proteins Etanercept and hGH,^3,4^ the majority of the soluble recombinant protein remained within the cell (Table 1). EM data show VNp-mNG-Etanercept impacts VNp remodelling of the inner membrane to induce VNp-fusion containing internalised cytosolic membrane structures (Fig. 2a-b). This IgG1 containing therapeutic fusion was not only dimeric (disulphide-bond dependent), but also exhibited appropriate ligand binding properties when isolated from VNp induced cytosolic vesicles, which was maintained upon TEV dependent proteolytic removal of the VNp-mNeongreen fusion (Supplementary Fig. 7). This is consistent with the lumen of the internal vesicle structures originating from the disulphide bond supporting inter-membrane periplasmic region.

**Figure 2.**
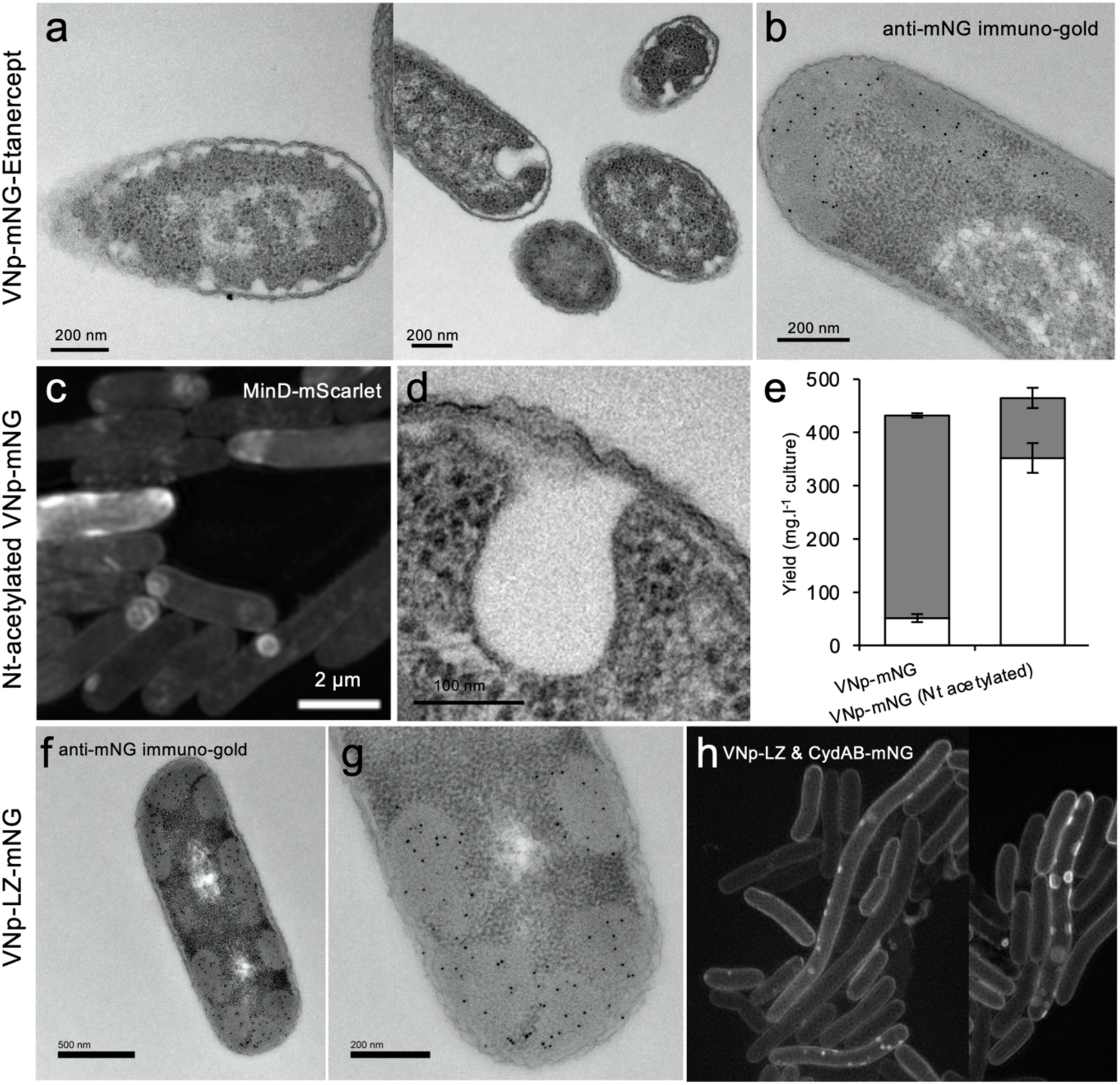
VNp dimer fusions produce fusion containing cellular membrane packages. (a) conventionally stained and (b) anti-mNeongreen immuno-stained EM serial section images of VNp-mNeongreen-Etancerpt induced inward membrane curvature in *E. coli*. (c) SIM imaging of mScarlet-MinD labelled inner membranes and (d) conventionally stained TEM sections of *E. coli* co-expressing VNp and the NatB amino-terminal acetylation complex producing cytosolic membrane bound structures. (e) Aminoterminal acetylation of VNp-mNeongreen promotes incorporation of the fusion protein within intracellular membrane bound structures (empty boxes) over export into extracellular vesicles (filled boxes). (f & g) Anti-mNeongreen immuno-EM images of sections though *E. coli* expressing VNp-LZ-mNeongreen, and (h) SIM images of CydAB^7^-mNeongreen labelled inner membranes in *E. coli* expressing VNp-LZ show the VNp-LZ dimer concentrates within the lumen of cytosolic inner membrane bound vesicles.

Similarly amino-terminal-acetylation of the VNp by recombinant NatB complex^5^ which, like VNp-Etanercept, brings about formation of internal membrane structures within *E. coli* (Fig. 2c-d), increasing the proportion of mNeongreen within the cell (Fig. 2e, Table 1). Stable alpha-helical VNp-dimers were created by introducing a leucine-zipper sequence^6^ between VNp and cargo (Supplementary Fig.8), and consistent with the previous dimeric VNp-fusions, induced formation of mNeongreen containing inner membrane associated vesicular structures within the bacterial cytosol (Fig. 2f-h; Supplementary Video 1). Therefore this innovation provides an attractive method for generating recombinant protein containing internal membrane bound structures for expression and compartmentalisation of disulphide bond containing and otherwise insoluble or toxic proteins from *E. coli*.

Co-expression of VNp-LZ dimeric cargo fusions with an additional VNp-LZ peptide (to increase overall extracellular vesicle production) resulted in the re-direction of internal compartment bound proteins towards the export route, as soluble cargo packaged vesicles isolated from the media (Table 1) to facilitate specific downstream processes for these proteins. Thus, not only does the VNp system support the immediate isolation of fusion proteins from the media, but it offers alternative internal expression systems, where this would be advantageous in certain fields such as the generation of enzyme cascades for complex synthesis or other aspects of synthetic biology.

Spurred on by the success of this approach, we asked whether simple modifications to the VNp amino acid sequence to modulate lipid interactions would enhance the exported protein yields. We therefore systematically tested a series of VNp variants, and by modifying charges and side chain length of targeted residues along the helix surface found that we could not only enhance vesicular export over a wide range of culture temperatures (VNp6), but also reduce the size of the VNp to 20 residues length (VNp15) to enhance the export of target model biopharmaceuticals at yields of more than 2.5 g of soluble recombinant protein / litre of bacterial flask culture (Table 1).

The VNp system exhibits a high degree of flexibility as vesicle packaged proteins can be generated in different *E. coli* cells (from a range of BL21 and K12 lineages, as well as DH10β and JM109 strains) making it perfectly suited for the production of synthetic proteins with modifications supported by specialist *E. coli* hosts. VNp-fusions can be expressed from a variety of plasmids (including pUC19 and pBR322 based derivatives) and modulated VNp-fusion expression can be supported from diverse promoters (e.g. T7, rhamnose, arabinose and Tac) and induction levels, making this a truly versatile system.

In summary, this simple peptide fusion increases yields and simplifies downstream processing of a wide range of recombinant proteins from *E. coli*. Importantly, the ease with which otherwise insoluble or toxic proteins can be isolated in milligram or gram quantities suggests that this approach is an attractive starting point for the expression of any recombinant protein of interest. A highly attractive aspect of this innovation is the stability of proteins and preservation of enzymatic activity when the vesicles are maintained at 4 ºC. As the use of this system is more broadly adopted and further enhancements and adaptations emerge, its impact can be anticipated to be highly significant. We therefore predict rapid adoption of this versatile system into a wide range of downstream processes and applications.

## Note

The Vesicle Nucleating peptide technology described here is associated with patent application *# GB2118435.3*.

## Supporting information

Eastwood et al - Supplementary data

