## Supplementary material for "Exceptional yield vesicle packaged recombinant protein production from *E. coli*": Eastwood et al - Supplementary data

### MATERIALS

#### *E. COLI* STRAINS USED IN THIS STUDY

| | |
|---|---|
| BL21 DE3 | $F^-$*ompT* *hsd*$S_B$ ($r_B^-$, $m_B^-$) *gal dcm* (DE3). |
| DH10β | $F^-$*mcr*A Δ(*mrr-hsd*RMS-*mcr*BC) φ80*lac*ZΔM15 Δ*lac*X74 *rec*A1 *end*A1 *ara*D139 Δ(*ara-leu*)7697 *gal*U *gal*K $\lambda^-$*rps*L($Str^R$) *nup*G |
| W3110 | $F^-$ $\lambda^-$ *IN(rrnD-rrnE)1 rph-1* |
| CLD1040 | $F^-$ *λ*, *IN(rrnD-rrnE)1 rph-1 OmpT* |
| JM109 | $F^-$ *tra*D36 *pro*AB *laq*$I^q$ZΔM15 *end*A1 *rec*A1 *gyr*A96 *thi hsd*R17 ($r_k^-$, $m_k^+$) *rel*A1 *sup*E44 Δ (*lac-pro*AB) |

### PLASMIDS

pRSFDuet-1_VNp-His6
pRSFDuet-1_VNp-mNeongreen
pRSFDuet-1_VNp6-mNeongreen
pRSFDuet-1_VNp15-mNeongreen
pRSFDuet-1_VNp-mNeongreen_OmpA-mCherry
pRSFDuet-1_VNp-mNeongreen_mScarlet-minD
pRSFDuet-1_VNp-mNeongreen_CydAB-mCherry
pETDuet-1_VNp-mCerulean3_Citrine-minD
pACYCDuet-1_*naa20_naa25* (pNatB) [1]
pRSFDuet-1_VNp-LZ-mNeongreen
pRSFDuet-1_VNp-LZ_CydAB-mNeongreen
pETDuet-1_VenusN154_VenusC155 (BiFC control construct)
pETDuet-1_VNp-VenusN154_VNp-VenusC155 (BiFC construct)
pETDuet-1_VNp-LZ-VenusN154_VNp-LZ-VenusC155 (BiFC construct)
pRSFDuet-1_DARPinOFF7
pRSFDuet-1_VNp-DARPinOFF7
pRSFDuet-1_VNp6-DARPinOFF7
pRSFDuet-1_VNp15-DARPinOFF7
pRSFDuet-1_Uricase
pRSFDuet-1_VNp-Uricase
pRSFDuet-1_StefinA
pRSFDuet-1_VNp-StefinA
pRSFDuet-1_VNp6-StefinA
pRSFDuet-1_VNp15-StefinA
pRSFDuet-1_FGF21
pRSFDuet-1_VNp-FGF21
pRSFDuet-1_DNAseI
pRSFDuet-1_VNp-DNAseI
pRSFDuet-1_hGH

pRSFDuet-1_VNp-hGH
pRSFDuet-1_VNp-LZ-hGH
pRSFDuet-1_mNeongreen
pRSFDuet-1_mNeongreen-DARPinOFF7
pRSFDuet-1_VNp-mNeongreen-DARPinOFF7
pRSFDuet-1_mNeongreen-Uricase
pRSFDuet-1_VNp-mNeongreen-Uricase
pRSFDuet-1_mNeongreen-StefinA
pRSFDuet-1_VNp-mNeongreen-StefinA
pRSFDuet-1_mNeongreen-Etanercept
pRSFDuet-1_VNp-mNeongreen-Etanercept
pRSFDuet-1_mNeongreen-Erythropoietin
pRSFDuet-1_VNp-mNeongreen-Erythropoietin
pRSFDuet-1_VNp-LZ_VNp-LZ-mNeongreen-Etanercept
pRSFDuet-1_VNp-LZ_VNp-LZ-mNeongreen-hGH

*Plasmid sequences and constructs will be available at addgene.org upon final publication.*

### POLYPEPTIDE SEQUENCES

(Residues differing from original VNp sequence are underlined).

**VNp peptide**: MDVFMKGLSKAKEGVVAAAEKTKQGVAEAAGKTKEGVL

**VNp6 peptide**: MDVFKKGFSIADEGVVGAVEKTDQGVTEAAEKTKEGVM

**VNp15 peptide**: MDVFKKGFSIADEGVVGAVE

**LZ peptide**: RMKQLEDKVEELLSKNYHLENEVARLKKLVG

Uniprot accession numbers of cargo full length protein sequences tested in this study:

- **DARP**- Designed Ankyrin Repeat Protein off7 (*Agrobacterium radiobacter*) - **B9JMD9**
- **DNaseI**- Deoxyribonuclease I (*Bos taurus*) - **P00639**
- **EPO** - Erythropoietin (*Homo sapiens*) - **P01588**
- **Etanercept** - Tumour necrosis factor receptor 1B - IgG1 fusion (*Homo sapiens*) - **P20333**
- **FGF21** - Fibroblast Growth Factor 21 (*Homo sapiens*) - **Q9NSA1**
- **hGH** - Somatotrophin (*Homo sapiens*) - **P01241**
- **mNeongreen** (*Branchiostoma lanceolatum*) - **A0A1S4NYF2**
- **StefinA** – Cystatin-A (*Homo sapiens*) - **P01040**
- **Uricase** (*Cyberlindnera jadinii*) - **P78609**

### METHODS

**Bacterial cell culture and protein induction**: All bacterial cells were cultured at 37 ºC in either LB (10 g Tryptone; 10g NaCl; 5g Yeast Extract (per litre)) or TB (12g Tryptone; 24g Yeast Extract; 4ml 10% glycerol; 17 mM $KH_2PO4$ 72 mM $K_2HPO_4$ (per litre)) media. 5 ml LB starters from fresh bacterial transformations were cultured at 37 ºC to saturation and used to inoculate 25 - 500 ml volume TB media, flask cultures that were incubated overnight at 37 ºC with 200 rpm and 25 mm throw orbital shaking. Recombinant protein expression from the T7 promoter was induced by addition of IPTG to a final concentration of 20 µg / ml (except Etanercept and hGH where 10 µg / ml was used) once the culture had reached an $OD_{600}$ of 0.8 – 1.0). To generate amino-terminally acetylated VNp, target constructs were co-transformed into *E. coli* with pNatB, to allow co-expression with the fission yeast amino-α-acetyltransferase complex B, Naa20 and Naa25 [1]. Amino-terminal acetylation was confirmed by electrospray mass spectroscopy of the purified VNp fusion protein. Growth curves were generated from 96 well plate cultures, prepared from late log-phase cultures, diluted into fresh media to an $OD_{600}$ of 0.1 at the start of the growth analysis experiment. $OD_{600}$ absorbance values were obtained using a Thermo Scientific Multiscan Go 1510-0318C plate reader and recorded using the SkanIt Software 4.0. at $OD_{600}$ values were taken every 30 minutes for the duration of the experiment, and growth curves generated from averages of ≥ 3 individual biological repeats.

**Soluble protein extracts**: Cell pellets from 25ml of a 50 ml culture were resuspended in 5 ml of soluble extract buffer (20 mM TRIS, 500 mM NaCl, pH 8.0), sonicated for a total of 2 min (6 x 20 sec pulses), and cell debris removed by centrifugation at 39,000 xg (4 ºC) for 20 min. Target protein concentration was determined using fluorescence of mNeonGreen fusion or gel densitometry. Both techniques were compared directly on the same samples to determine equivalence.

**Recombinant vesicle isolation**: Vesicles were isolated directly from overnight bacterial cell cultures by passing the culture through a sterile and detergent-free 0.45 µm polyethersulfone (PES) filter. Typical purity and concentration from equivalent volume of culture and filter flow through are shown in Fig. 1. Exclusion of viable cells from the vesicle containing filtrate was tested by plating onto LB plates lacking antibiotics and incubating overnight at 37 ºC (example shown in Fig. S4).

**Protein concentration determination**: Fluorescence scanning was used to determine concentration of mNeongreen fusion proteins in vesicle containing media and soluble protein extracts. Absorbance was measured at 506 nm using a Varian Cary 50 Bio UV-Vis spectrophotometer, with measurements from conditioned media used for baseline correction, and concentration determined using an extinction coefficient of 116,000 $M^{-1}cm^{-1}$. Concentration of non-mNeongreen labelled proteins was determined by gel densitometry analysis of triplicate samples run alongside BSA loading standards on coomassie stained SDS-PAGE gels. Gels were scanned and analysed using Image J software. Protein concentrations were determined by both UV and densitometry for three independent VNp-mNeongreen samples to confirm parity between analysis techniques. Average yields in Fig. 1 & Table S1 were calculated (mg soluble target protein / litre culture) from a minimum of 3 independent biological repeats from cultures of BL21 DE3 *E. coli* cells grown in TB media.

**Determination of cytosolic VNp concentration**: VNp-mNeongreen expression was induced in BL21 DE3 *E. coli* for 4 hours at 37ºC (when extracellular vesicle production is observed), and images of more than 120 cells were acquired using widefield imaging (described below) from 3 independent sample preparations. Mean mNeongreen intensity was determine from a 5 x 5 pixel area within each *E. coli* cell. A calibration line was generated from multiple images of known concentrations of VNp-mNeongreen solutions using identical imaging conditions as cell image acquisition. Using these data, the average cytosolic VNp-mNeongreen concentration at the point of vesicle production was calculated to be 18.76 ± 0.14 µM.

**Affinity purification of His-tagged proteins**: To isolate recombinant protein from vesicles, after a 10 min 3,000g centrifugation to pellet cells, media from an induced overnight culture expressing VNp-labelled protein was passed through a 0.45 µm PES filter, and the subsequent vesicle containing flow-through was sonicated and mixed in a 1 in 5 dilution of 5 x binding buffer (250 mM TRIS 2.5 M NaCl 5% Triton-X 50 mM Imidazole pH 7.8) before passing over a $Ni^{2+}$-agarose resin gravity column. Cytosolic recombinant protein was purified by passing soluble protein extracts (supplemented with Imidazole to 20 mM) over the $Ni^{2+}$-agarose resin gravity column. In both cases matrix bound His-tagged protein was washed, eluted (using imidazole), and dialysed into appropriate storage or assay buffer. Protein identity and amino-terminal acetylation of isolated proteins was confirmed by electrospray mass-spectroscopy.

**Circular Dichroism (CD)**: Measurements were made in 2 mm quartz cuvettes using a Jasco 715 spectropolarimeter. VNp protein and 100 nm extruded vesicles were diluted in CD buffer (10 mM potassium phosphate, 5 mM $MgCl_2$ pH 7.0) to a concentration of 0.4 mg/ml and 0.2 respectively. Broad negative peaks at 208 and 222 nm and a positive peak at < 200 nm are consistent with an α-helical structure.

**Dynamic Light Scattering studies:** All studies were carried out using Anton Paar Litesizer$^{TM}$ 500 and processed using KalliopeTM Professional. All vials/cuvettes/microfuge tubes used for sample preparation were clean and dry. All solvent systems used were filtered to remove any particulates that may interfere with the results obtained.

**Electrospray LC-MS of Proteins**: Electrospray mass spectra were recorded on a Bruker micrOTOF-Q II mass spectrometer. Samples were desalted on-line by reverse-phase HPLC on a Phenomenex Jupiter C4 column (5 µm, 300 Å, 2.0 mm x 50 mm) running on an Agilent 1100 HPLC system at a flow rate of 0.2 ml/min using a short water, acetonitrile, 0.05% trifluoroacetic acid gradient. The eluant was monitored at 214 nm & 280 nm and then directed into the electrospray source, operating in positive ion mode, at 4.5 kV and mass spectra recorded from 500-3000 m/z. Data was analysed and deconvoluted to give uncharged protein masses with Bruker's Compass Data Analysis software.

**Gel filtration assay:** 500 µl of protein samples were loaded to a Superdex 200 Increase 10/300 GL size-exclusion column (GE Healthcare Life Sciences) equilibrated at room temperature in PBS and run at 0.75 ml/min flow rate. Eluted proteins were measured by Viscotek Sec-Mals 9 and Viscotek RI detector VE3580 (Malvern Panalytical). Data was analysed using OmniSEC software.

**Lipid binding Assay:** Affinity of VNp for *E. coli* membrane lipids was established using a thermal shift fluorescence binding assay adapted from [2]. Equivalent assay samples, comprising: 65 µl 3 mg/ml of VNp-mNeongreen, 65 µl 1 mM of 100 nm extruded vesicles composed of the lipid mixture to be tested; 15 µl, 10%OGP; and 5 µl 20 mM Tris-HCl pH 7.0, were prepared in 0.3 ml microfuge tubes and held at the defined temperature in a gradient Techne PCR machine for 10 minutes. Samples were centrifuged at 18,000 *xg*, and supernatant fluorescence was determined in black 96 well plates (BRAND, Germany) using a BMG Clariostar (BMG Labtech). Fluorescence readings were normalised and used to create a melting curve, where the melting temperature (*Tm*) was determined using Origin software (OriginLab). The final *Tm* value was an average (± s.d) calculated from three independent sample repeats.

**Uricase Assay**: Uricase activity was either measured using a Varian Cary 50 Bio UV-Vis spectrophotometer (cuvette measurement) or using a TgK HiTech stopped flow system. For the cuvette measurements 500 µl of 100 mM Tris pH 8.5 with 200 mM Uric acid was placed in a cuvette and $OD_{293}$ measurements were taken over for 4 or 5 minutes. Subsequently either 500 µl of 4.5 mg/ml purified VNp2-Uricase (dialysed into 0.1 M Tris ph8.5) or dialysis buffer alone was added to the cuvette and $OD_{293}$ measurements taken for 25 min. (Adapted from [3]). For stopped flow method, absorbance was measured at 290 nm at 20 °C using a 22.5 µl observation cell, and the reaction was followed for 2,000 seconds. Rates were determined from linear fits to the steady state region of a curve average from 3 independent reactions.

**Widefield Fluorescence Microscopy**: Cells were mounted onto coverslips under < 1 mm thick circular LB-agarose(2%) pads, and attached with appropriate spacers onto glass slides, before being visualised on an inverted microscope [4]. Live cell imaging for each sample was completed within 30 min of mounting the cell sample onto coverslips.

**Structured Illumination Microscopy (SIM)** was undertaken using a Zeiss Elyra PS 1 microscope with a 100x NA 1.46 oil immersion objective lens (Zeiss α Plan-Apochromat) as described previously.[5,6] Briefly, cells were mounted under thin LB-agarose pads onto high precision No.1.5 coverslips (Zeiss, Jenna, Germany). 488 nm and 561 nm laser were used to illuminate mNeongreen and mCherry/mScarlet fusions, respectively. The optical filter set consisted of laser blocking filter MBS 405/488/561 (Chroma) as the dichroic mirror, and the dual-band emission filter LBF-488/561. The total of 3 rotations of the illumination pattern were implemented to obtain two-dimensional information. Super-resolution SIM image processing was performed using the Zeiss Zen software. Two colour images were aligned using the same software following a calibration using pre-mounted MultiSpec bead sample.

**Fluorescence Lifetime Imaging Microscopy (FLIM):** The one- and two- photon systems used in this work have been previously described [7]. Prior to FLIM data acquisition, protein expression levels were verified using confocal microscope. Here, a Nikon Eclipse C2-Si confocal scan head attached to an inverted Nikon TE2000 or Ti-E microscope was used. mNeongreen and mCherry FP were excited at 491 nm (emission 520/35 nm) and 561 nm (emission 630/50 nm) respectively using an NKT super continuum laser. FLIM images were obtained as follows: 2 photon (950 nm) wavelength light was generated by a mode-locked titanium sapphire laser (Mira F900, Coherent Laser Ltd.), producing 180 fs pulses at 76 MHz. This laser was pumped by a solid-state continuous wave 532 nm laser (Verdi 18, Coherent Lasers Ltd.). Fluorescence was collected through a BG39 filter for the donor fluorophore. The acceptor was not excited.
For one photon excitation FLIM, the system is equipped with a SuperK EXTREME NKT-SC 470-2000 nm supercontinuum laser (NKT Photonics) which generates at 80 MHz repetition rate with 70 ps pulse width. The desired wavelengths were selected using a SuperK SELECT 29 multi-line tunable filter (NKT photonics). Images were collected through either a 60X 1.2 NA water immersion (Fig S1c) or 60X 1.49 NA oil immersion (Fig S1d & e) objective lens. For both one and two-photon excitation, emission was collected by the same objective through filters (above) and detected with an external hybrid GaAsP (HPM-100-40, Becker & Hickl, Germany), linked to a time correlated single photon counting (TCSPC) module (SPC830, Becker and Hickl, Germany). Photon counts of at least 1000 used for the multi-exponential analysis. Raw time correlated single photon counting decay curve at each pixel (256 x 256 or higher) of the images were analysed using SPCImage software v.6.9 (Becker and Hickl, GmbH); an incomplete single exponential fit model with a laser repetition time value of 12.5 ns was used for the decay curve fitting. Lifetime values with $\chi 2$ between 0.8 and 1.3 were taken as a good exponential decay fit. Lifetimes were calculated from an average of a minimum of 15 distinct fields of view, with ≥ 5 taken from 3 separately prepared slide samples.

**Transmisssion Electron Microscopy (TEM) analysis of cells and isolated vesicles**: Negative stained TEM samples of cells and vesicles were prepared in one of two ways.10 µl of *E. coli* cells expressing VNp-mNeongreen from an overnight culture was placed onto a formvar/carbon coated 400mesh gold grid and incubated in a humid chamber at 37 ºC to allow vesicle formation. Recombinant vesicles isolated from a culture of *E. coli* expressing VNp-mNeongreen were placed onto a formvar/carbon coated 600-mesh copper grid and left for 5 min at room temperature to allow vesicles to settle onto the surface. Both samples were then fixed in 2.5% glutaraldehyde in 100 mM sodium cacodylate buffer pH 7.2 (CAB) for 10 minutes. Grids were then washed in 100 mM CAB and milliQ water. Grids were then dried and negative stained for 5 seconds in 2% aqueous uranyl acetate.

**TEM thin section analysis of *E. coli* cells**: *E. coli* expressing VNp-mNeongreen (± fusions) were cultured as described above and harvested by centrifugation at 3,000 *g* for 10 min. The cell pellet (approximately 100µl) was resuspended in 2 ml of 2.5% (w/v) glutaraldehyde in CAB and fixed for 2 hr at room temperature with gentle rotating (20 rpm). Cells were pelleted by centrifugation at 6,000 *g* for 2 min and were washed twice for 10 min with 100 mM CAB. Cells were postfixed with 1% (w/v) osmium tetroxide in 100 mM CAB for 2 hr and subsequently washed twice with ddH2O. Cells were dehydrated by incubation in an ethanol gradient, 50% EtOH for 10 min, 70% EtOH overnight, and 90% EtOH for 10 min followed by three 10 min washes in 100% dry EtOH. Cells were then washed twice with propylene oxide for 15 min. Cell pellets were embedded by re- suspension in 1 ml of a 1:1 mix of propylene oxide and Agar LV Resin and incubated for 30 min with rotation. Cell pellets were infiltrated twice in 100% Agar LV resin (2 x 2h). The cell pellet was resuspended in fresh resin and transferred to a 1-mL BEEM embedding capsule, centrifuged for 5 min at 1100 rpm in a swing out rotor to concentrate the cells in the tip of the capsule and samples were polymerised for 20 hr at 60°C.

Ultrathin sections were cut using a Leica EM UC7 ultramicrotome equipped with a diamond knife (DiATOME 45°). Sections (70 nm) were collected on uncoated 400-mesh copper grids.

Grids were stained by incubation in 4.5% (w/v) uranyl acetate in 1% (v/v) acetic acid for 45 min followed by washing in a stream of ddH2O. Grids were then stained with Reynolds lead citrate for 7 min followed by washing in a stream of ddH2O. Electron microscopy was performed using a JEOL-1230 transmission electron microscope operated at an accelerating voltage of 80 kV equipped with a Gatan One View digital camera.

**Immuno-EM of isolated vesicles:** 2µl of filtered media containing recombinant vesicles from a culture of *E. coli* expressing VNp-mNeongreen was placed onto a formvar/carbon coated 600-mesh copper grid and left for 5 min at room temperature to allow vesicles to settle. Vesicles were osmotically shocked to rupture vesicles by moving grids into 2 x 20µl drops of milliQ water for 10 minutes at RT. Samples were then fixed in 2% formaldehyde and 0.5% EM grade glutaraldehyde in CAB for 15 minutes at RT. Grids were then washed in 6 x 20µl drops of CAB and 6 x 20µl drops of TBST (20 mM Tris-HCl, 500 mM NaCl, 0.05% Tween 20 and 0.1% BSA pH7.4). Samples were blocked in a 20µl drop of 2% BSA in TBST at room temperature for 30 min. Grids were then transferred directly into a 20µl drop of anti-mNeongreen rabbit polyclonal (Cell Signalling Technology) primary antibody diluted 1:100 in TBST and incubated for 1 hr. Grids were washed in 6 x 20µl drops of TBST. Grids were then moved into a drop of goat anti-rabbit IgG 5nm gold (British Biocell International) diluted 1:50 and then moved to a fresh drop of the same antibody and incubated for 30 min. Excess antibody was removed by washing in 6 x 20µl drops of TBST and 6 x 20µl drops of milliQ water and dried.

Grids were negative stained for 5 seconds in 2% aqueous uranyl acetate. Electron microscopy was performed using a JEOL-1230 transmission electron microscope operated at an accelerating voltage of 80 kV equipped with a Gatan One View digital camera.

**Immuno-EM of *E. coli* cells:** *E. coli* expressing VNp-mNeongreen were cultured as described above and harvested by centrifugation at 3,000 *g* for 10 min. The cell pellet (approximately 100µl) was resuspended in 2 ml 2% (w/v) formaldehyde and 0.5% glutaraldehyde in CAB and fixed for 2h at RT. The sample was washed 2 x10 min in CAB. Cells were dehydrated by incubation in an ethanol gradient, 50% EtOH for 10 min, 70% EtOH overnight, and 90% EtOH for 10 min followed by three 10 min washes in 100% dry EtOH. Cells were then suspended in LR White resin medium grade (London Resin Company) for 4 hr and then in fresh LR White resin overnight. Following 2 x 4 hr changes in fresh LR White resin samples were placed in sealed gelatine capsules and spun in a swing out rotor at 1,100rpm to concentrate cells. Gelatine capsules containing the cell pellets were polymerised upright at 60 ºC for 20 hr. Ultrathin sections were cut using a Leica EM UC7 ultramicrotome equipped with a diamond knife (DiATOME 45°). Sections (80 nm) were collected on uncoated 400-mesh gold grids.

Samples were blocked in a 20µl drop of 2% BSA in TBST at room temperature for 30 min. Grids were then transferred directly into a 20µl drop of anti-mNeongreen rabbit polyclonal (Cell Signalling Technology) primary antibody diluted 1:10 in TBST and incubated for 1 hr. Grids were washed in 6 x TBST. Grids were then moved into a drop of goat anti-rabbit IgG 5nm gold (British Biocell International) diluted 1:50 and then moved to a fresh drop of the same antibody and incubated for 30 min. Excess antibody was removed by washing in 6 x 20µl drops of TBST and 6 x 20 µl drops of milliQ water and dried.

Grids were stained for 15 min in 4.5% uranyl acetate in 1% acetic acid solution and then washed in 6 x 20µl drops of milliQ water. Grids were then stained with Reynolds lead citrate for 3 min and washed in 6 x 20µl drops of milliQ water. Electron microscopy was performed using a JEOL-1230 transmission electron microscope operated at an accelerating voltage of 80 kV equipped with a Gatan One View digital camera.

### ACKNOWLEDGEMENTS:

The authors thank J. Walklate for assistance undertaking stopped-flow experiments, S. Boxall, M. Geeves, I. Hagan, and D. Manstein for stimulating discussions and comments on the manuscript. This work was supported by the University of Kent and funding from the Biotechnology and Biological Sciences Research Council (BB/S005544/1 & BB/L013703/1_D0101) and Fujifilm-Diosynth Biotechnologies UK Ltd.

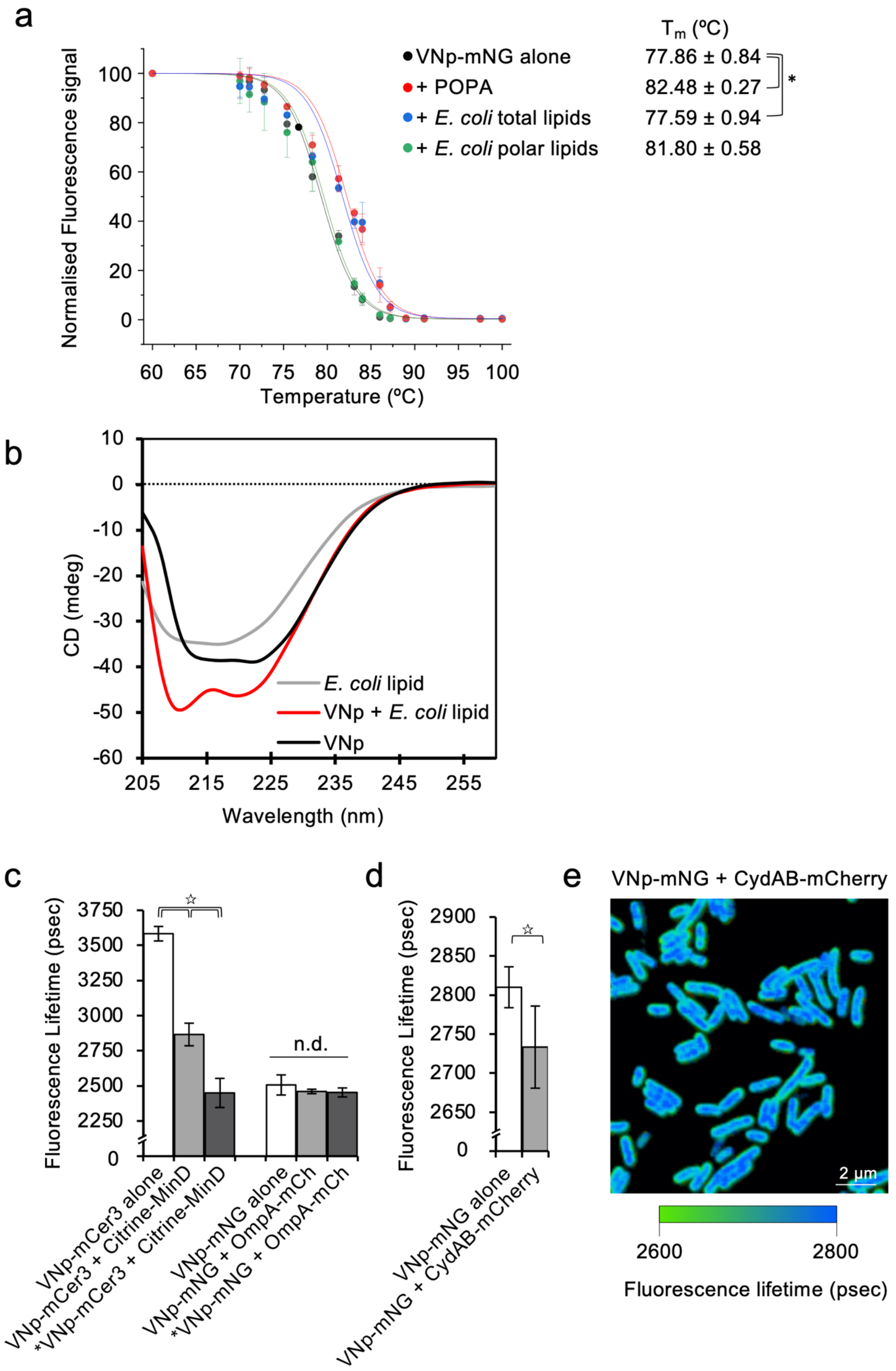
a
Tm (ºC)
VNp-mNG alone 77.86 ± 0.84
+ POPA 82.48 ± 0.27
+ E. coli total lipids 77.59 ± 0.94
+ E. coli polar lipids 81.80 ± 0.58
*
Normalised Fluorescence signal
Temperature (ºC)
b
CD (mdeg)
Wavelength (nm)
E. coli lipid
VNp + E. coli lipid
VNp
c
Fluorescence Lifetime (psec)
n.d.
VNp-mCer3 alone
VNp-mCer3 + Citrine-MinD
*VNp-mCer3 + Citrine-MinD
VNp-mNG alone
VNp-mNG + OmpA-mCh
*VNp-mNG + OmpA-mCh
d
Fluorescence Lifetime (psec)
VNp-mNG alone
VNp-mNG + CydAB-mCherry
e
VNp-mNG + CydAB-mCherry
2 µm
2600
2800
Fluorescence lifetime (psec)

**Supplemental Figure 1. VNp interacts with the inner *E. coli* membrane.** Thermal shift (a) and Circular Dichroism (b) assays were used to confirm interaction between the VNp and membrane composed of *E. coli* membrane lipids *in vitro.* (a) Interaction with the membrane increases thermal stability of a membrane associated fluorophore. The thermal shift assay was used to examine the impact of different lipid membranes upon fluorophore signal from a fluorescent protein (mNeongreen) when fused to the VNp, a potential membrane binding protein. Average Thermal shift mNeongreen fluorescence curves were calculated for VNp-mNeongreen alone (black), and VNp-mNeongreen in the presence of 100 nm vesicles composed of phosphatidic acid (red), or mixtures of either total (blue) or polar (green) *E.coli* lipids. The shift to the right signifies the VNp-mNeongreen interacts with membranes composed of phosphatidic acid or a mixture of total *E. coli* lipids. (b) Circular Dichroism was used to examine the impact *E. coli* membrane binding has upon the predicted alpha-helical VNp structure. The graph shows averaged CD spectra of VNp alone (black), total *E. coli* lipid vesicles alone (grey), or from a mixture of VNp and *E. coli* lipid vesicles (red). The relative broad negative CD spectra peaks at 208 nm and 222 nm, observed in the mixture of VNp and *E. coli* lipid membrane are consistent with single α-helical structures, and these spectra show that the VNp alpha-helix is stabilised upon interaction with *E. coli* membrane lipid vesicles. (c-e) Fluorescence Lifetime Imaging based Fluorescence Resonance Energy Transfer Microscopy (FLIM-FRET) was used to examine physical interactions between VNp and the *E. coli* inner and outer membranes *in vivo*. Cerulean3 (VNp-Cer3) or mNeongreen (VNp-mNG) fluoreophores were used as donors, and Citrine (Citrine-MinD) or mCherry (OmpA-mCherry/CydAB-mCherry) were used as acceptors. FRET dependent reduction in the fluorescence lifetime of the donor indicates physical interaction (< 10 nm) between proteins (☆ - 99.99% confidence levels). (c) Histogram of donor fluorophore Fluorescence lifetimes of *E. coli* cells expressing VNp donor fluorophore fusions (VNp-Cer3 / VNp-mNG) either alone or with acceptor fluorophore labelled inner (Citrine-MinD) or outer (OmpA-mCherry) membrane proteins indicate VNp interacts with the *E. coli* inner membrane. (d) The histogram of mNeongreen fluorescence lifetime within *E. coli* cells expressing an mNeongreen donor fluorophore VNp fusion (VNp-mNG alone or in combination with the mCherry labelled CydAB inner membrane complex confirmed interaction between VNp and the inner membrane. (e) mNeongreen fluorescence Lifetime micrograph of VNp-mNG CydAB-mCherry expressing cells (from d) illustrates the reduced fluorescence lifetime of the VNp-mNeongreen at the cell membrane, where CydAB is located. The reduction in lifetime length reflected in change from blue (2.8 ns) to green (2.6 ns).

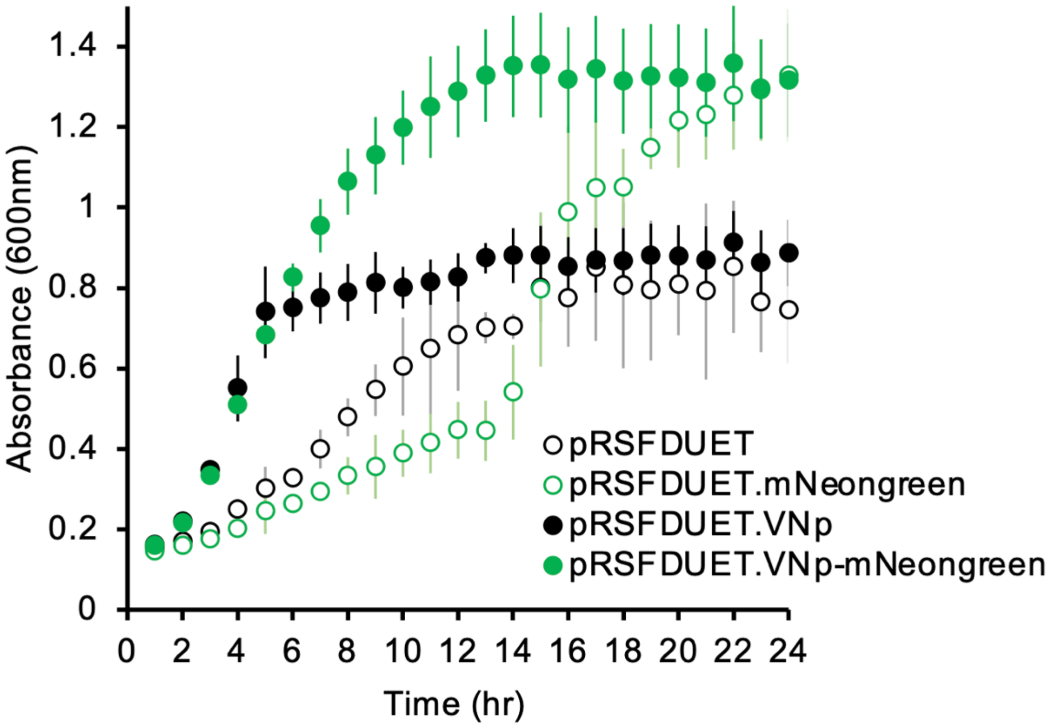


**SUPPLEMENTAL FIGURE 2: VNp expression does not impact *E. coli* growth or viability.** To assess the impact of VNp expression on *E. coli* viability averaged growth curves were generated from 4 independent replicate cultures of BL21(DE3) *E. coli* cells containing either an empty pRSFDUET vector (empty black circles), pRSFDUET.mNeongreen (empty green circles), pRSFDUET.VNp (filled black circles) or pRSFDUET.VNp.mNeongreen (filled green circles). Cells were grown at 37 ºC in TB supplemented with kanamycin and 20 µg/ml IPTG on the same 96-well plate. These data illustrate expression of VNp or a VNp fusion does not negatively impact bacterial growth or viability over a 24 hour period.

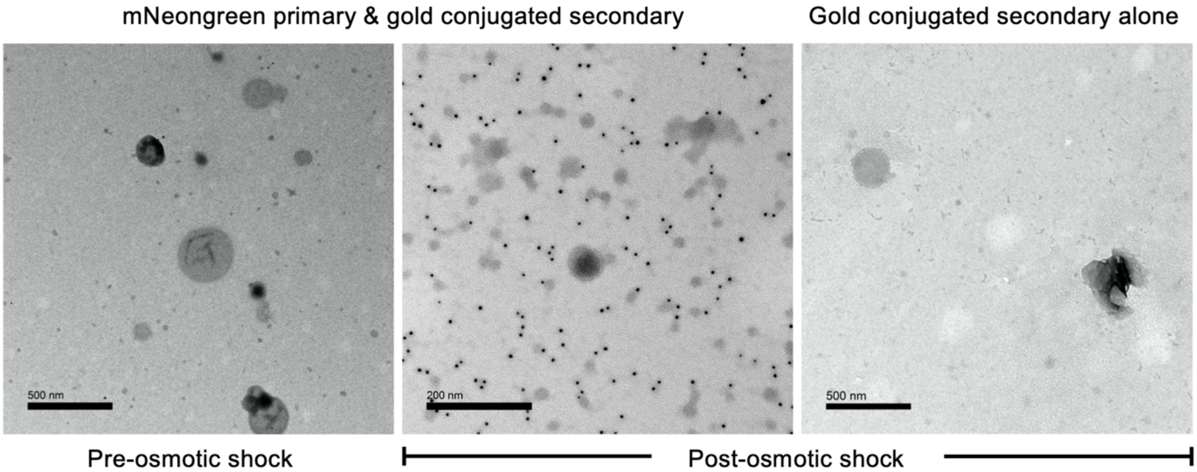


**SUPPLEMENTAL FIGURE 3: VNp fusion protein is contained within the lumen of isolated recombinant vesicles.** VNp-mNeongreen containing vesicles were filter purified from media of an overnight culture of BL21 DE3 pRSFDUET1-VNp-mNeongreen cells and mounted onto EM grids and subjected to anti-mNeongreen immuno EM analysis. VNp-mNeongreen dependent gold labelled densities bound to mNeongreen released from vesicles upon osmotic shock from resuspension in water. The lack of densities in control samples subjected to either immuno-analysis prior to bursting, or burst vesicles processed in the same way but without primary anti-mNeongreen antibodies illustrate the mNeongreen is located exclusively within the lumen of the vesicles.

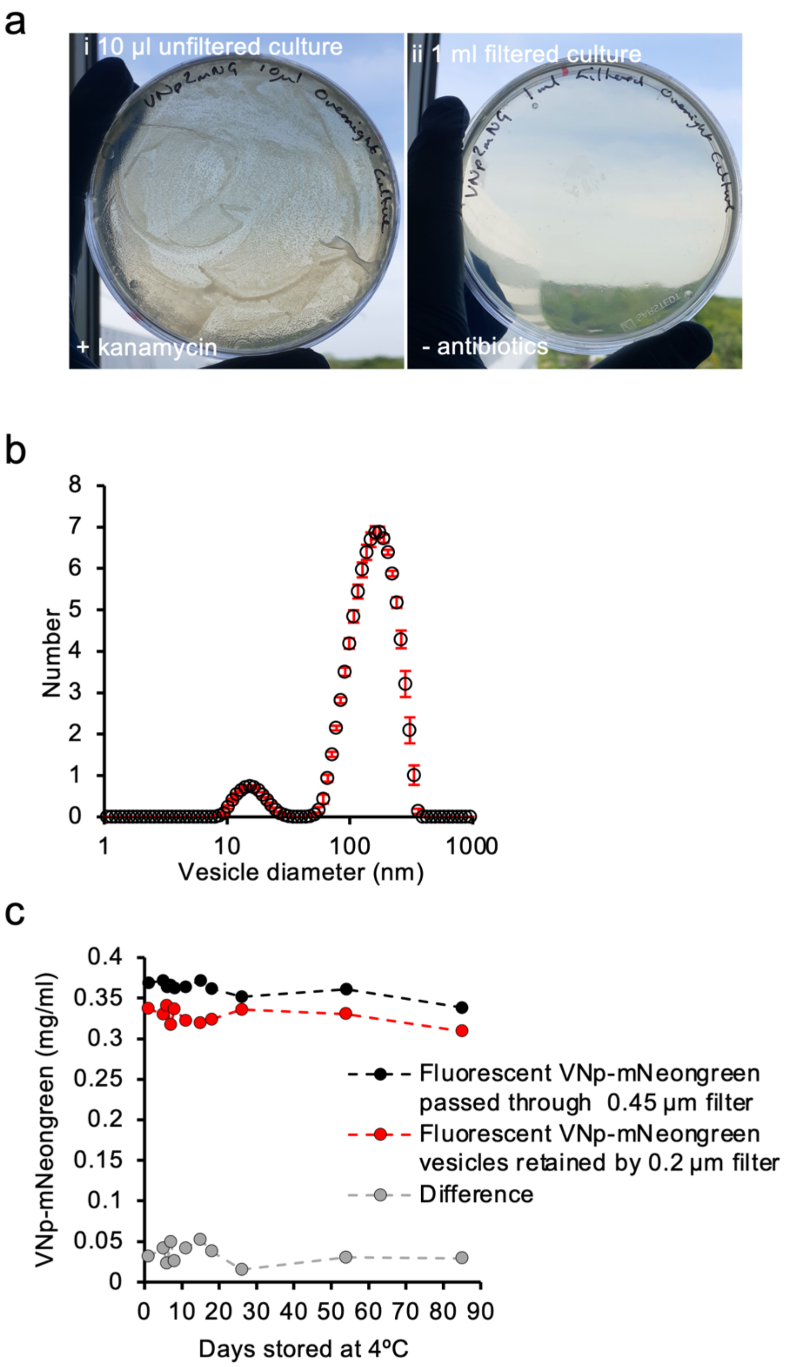


**Supplemental Figure 4: Filter purified VNp vesicles are bacteria free, uniform in size, and provide a stable environment for storage of VNp-fusions.** (a) Test illustrating exclusion of viable *E. coli* cells from the vesicle containing filtrate. 10 µl of total culture (i) and 1 ml of 0.45µm media filtrate (ii) from an overnight culture of VNp-mNG expressing *E. coli* cells were plated out onto LB (ii) or LB supplemented with kanamycin (i) and incubated overnight at 37 ºC. (b) Dynamic light scattering was used to examine the size of isolated VNp induced vesicles. The size profile of a population of filter purified VNp-mNeongreen induced vesicles illustrate the vesicles have a uniform size of 166 nm. The graph presents averaged data (± standard errors) from 10 separate runs. (c) The VNp induced vesicle provised a stable environment for storage of VNp-fusions. VNp-mNeongreen containing vesicles, filter purified from media of an overnight culture of BL21 DE3 pRSFDUET1-VNp-mNeongreen cells were stored at 4 ºC. Overall mNeongreen fluoresecence and the fraction of mNeongreen fluorescence within vesicles retained by a 0.2µm filter did not vary over time, indicating stability of vesicles and folded mNeongreen protein within them.

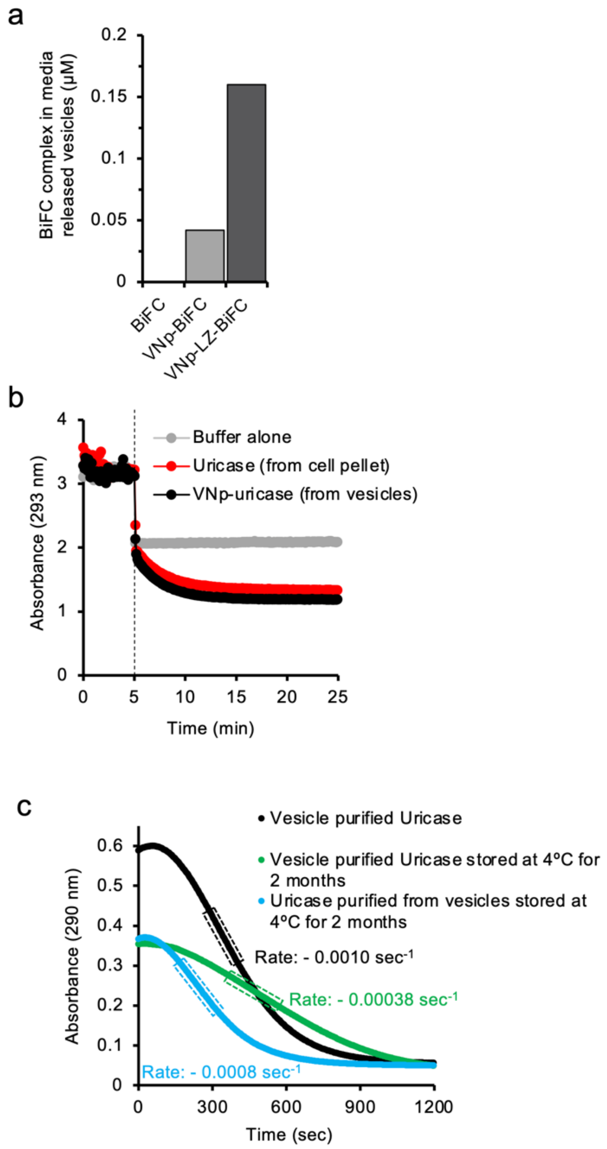


**SUPPLEMENTAL FIGURE 5: Heterodimeric and functional VNp-fusions are targeted to the VNp-vesicles.** (a) Bimolecular Fluorescence Complementation dependent Fluorescence between the amino (VenusN154) and carboxyl (VenusC155) was used to establish heterocomplexes between different VNp fusions could be targeted to and isolated from the same vesicle. VenusN154-VenusC155 BiFC dependent fluorescence (515 nm) from vesicles isolated from BL21(DE3) *E. coli* expressing either VenusN154 & VenusC155 (white); VNp-VenusN154 & VNp-VenusC155 (grey); or VNp-LZ-VenusN154_VNp-LZ-VenusC155 (black) was used to establish the concentration of the BiFC complex within isolated vesicles. (b) The enzymatic activity of Uricase either isolated from a cell pellet using conventional methods (red) or from VNp-uricase induced vesicles (black) was examined to compare functionality of each protein. A buffer only control (grey) shows dilution dependent change in baseline 293 nm absorbance of uric acid. Uricase enzyme / buffer was added to the uric acid substrate after a 5 min equilibration (dashed line). The activity of uricase isolated from cell pellet or VNp-uricase containing vesicles were equivalent. (c) To examine the stability of uricase enzyme activity from protein stored within VNp-uricase vesicles, uricase activity of either fresh vesicle purified VNp-uricase (black; the same vesicle purified VNp-uricase stored in reaction buffer at 4 ºC for 2 months (green); or freshly purified from vesicles that had been stored at 4 ºC for 2 months (blue), were measured using stopped-flow. Rates were determined from steady state regions (highlighted by boxes) of averaged curves and show while uricase stored in buffer exhibited 38% of the original activity, uricase stored within vesicles retained 80% of the original enzymatic activity.

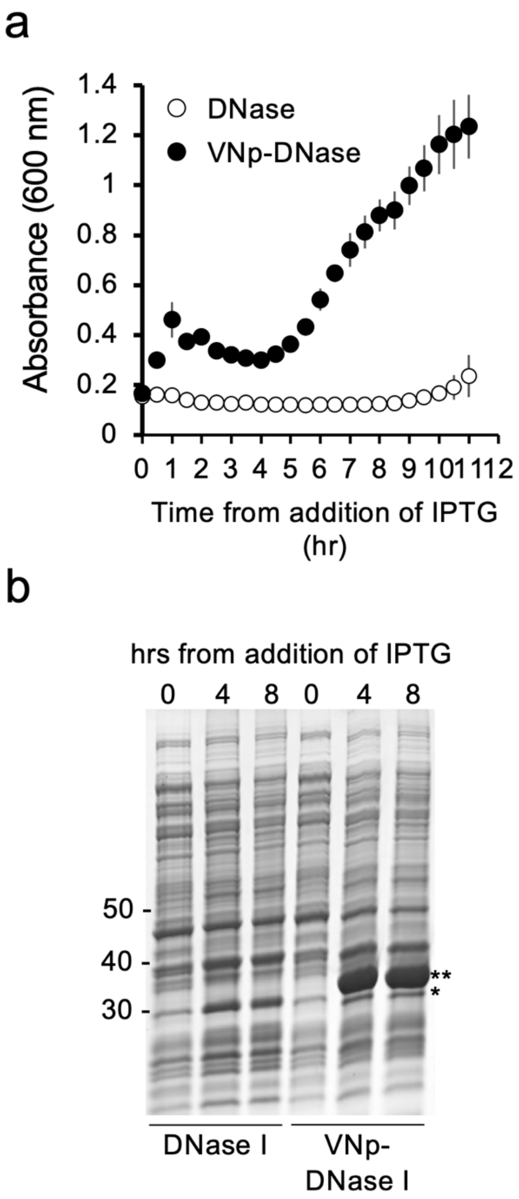


**SUPPLEMENTAL FIGURE 6: A VNp fusion permits expression of DNAseI in *E. coli*.** To establish whether vesicular compartmentalisation of VNp fusions allowed expression of toxic proteins the expression and impact on *E. coli* growth of DNAse1 and VNp-DNaseI were compared. (a) Average growth curves and (b) expression profiles of *E. coli* expressing DNase and VNp-DNase show that while DNaseI only had minimal expression, it inhibited growth of the *E. coli* cells. In contrast the VNp-DNAseI expressed cells grew normally and expressed meaningful levels of the fusion protein.Predicted sizes of DNase and VNp-DNase are 30.2 and 34.4 kDa respectively.

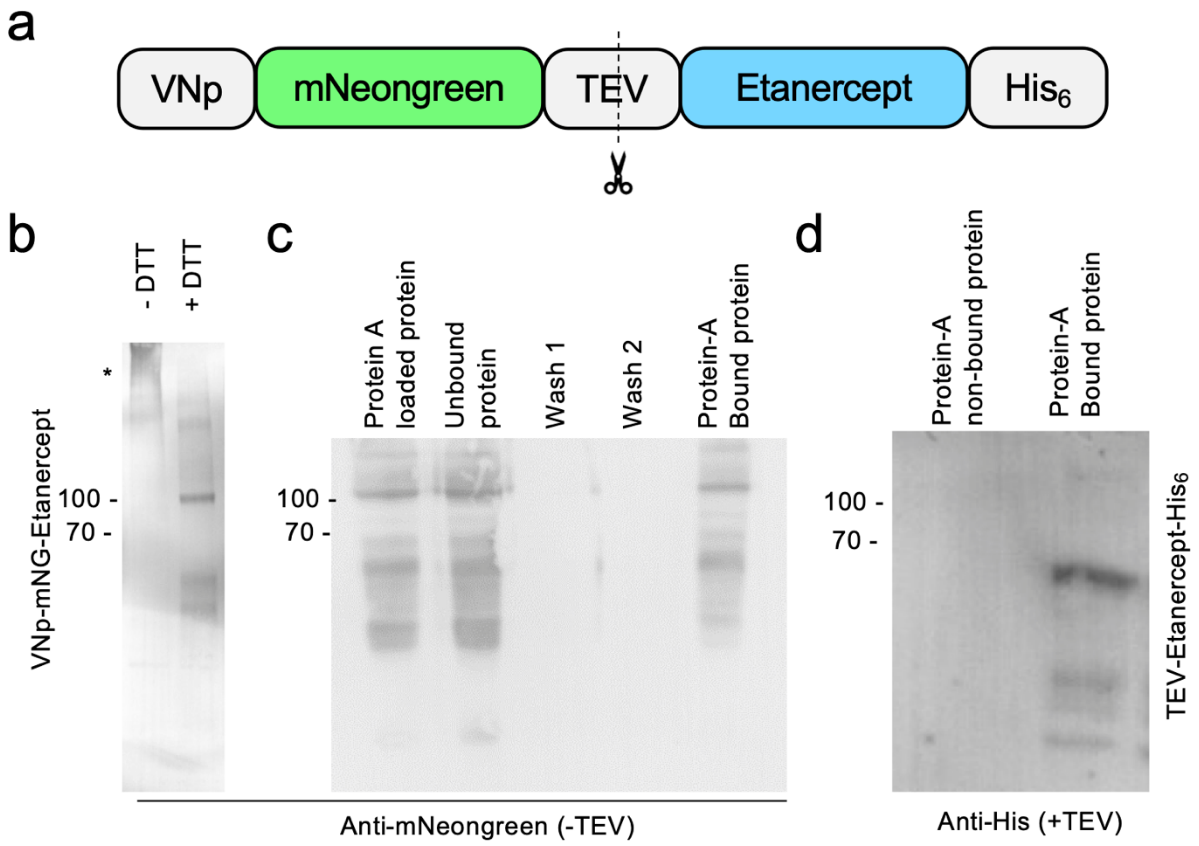


**SUPPLEMENTAL FIGURE 7: VNp allows expression of functional disulphide bond containing IgG1 fusion dimers.** VNp dependent targeting to the periplasm and subsequent cytosolic vesicles facilitates disulphide bond formation and functionality of the chimeric IgG1 containing fusion protein, Etanercept. (a) Schematic of the VNp-mNeongreen-TEV-Etanercept fusion protein. (b & c) Anti-mNeongreen western blots illustrating disulphide bond dependent oligomerisation (b) and protein A binding IgG1 functionality (c) of VNp-mNG-Etanercept purified from *E. coli*. (b) VNp-mNG-Etanercept disulphide bond-dependent oligomers (*) are disrupted by the addition of the disulphide bond disrupting reducing agent, DTT. (c) VNp-mNG-Etanercept-$His_6$ fusion was affinity purified from *E. coli* and bound to Protein A – Dynabeads. These beads were subsequently washed in binding buffer, before being boiled in SDS-PAGE loading buffer to release bound proteins. Predicted size of VNp-mNeongreen-Etanercept: 83.9 kDa. (d) Anti-His western blot of wash (unbound) and Protein A bound fractions of TEV cleaved VNp-mNeongreen-TEV-Etanercept-His fusion mixed with Protein A-Dynabeads. This illustrates unlabelled Etanercept remains soluble and functional.

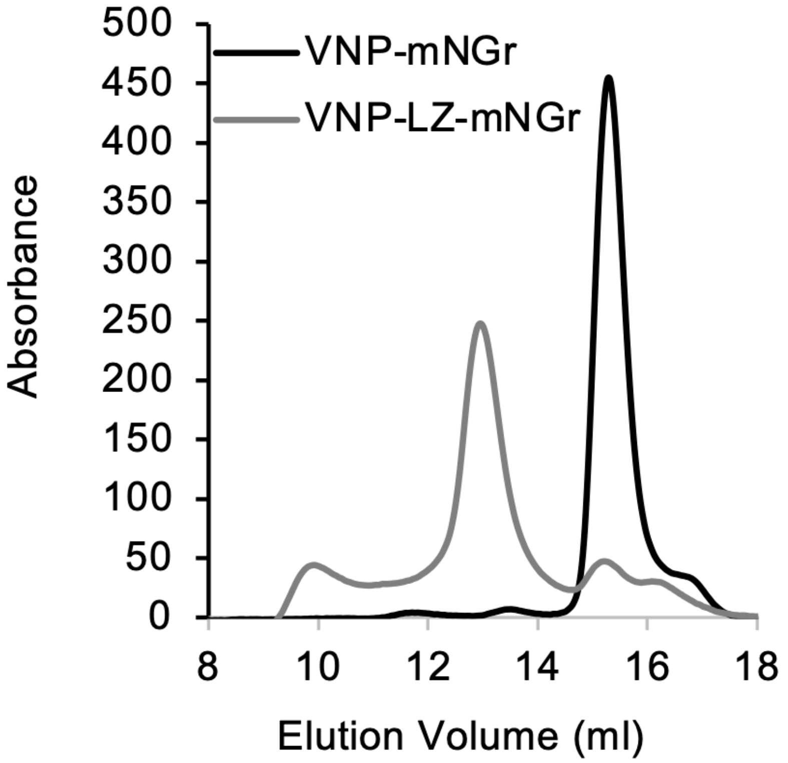


**SUPPLEMENTAL FIGURE 8: A VNp-Leucine Zipper fusion is dimeric.**
Gel filtration analysis was undertaken to confirm whether introducing a Leucine Zipper (LZ) motif to the VNp-mNeongreen (mNGr) fusion brought about stable dimers formation. While the elution profile of VNp-mNGr (black) were consistent with a monomeric protein, the VNp-LZ-mNG (grey) eluted from the column within earlier fractions, consistent with dimeric status of the VNp-LZ-mNGr fusion.

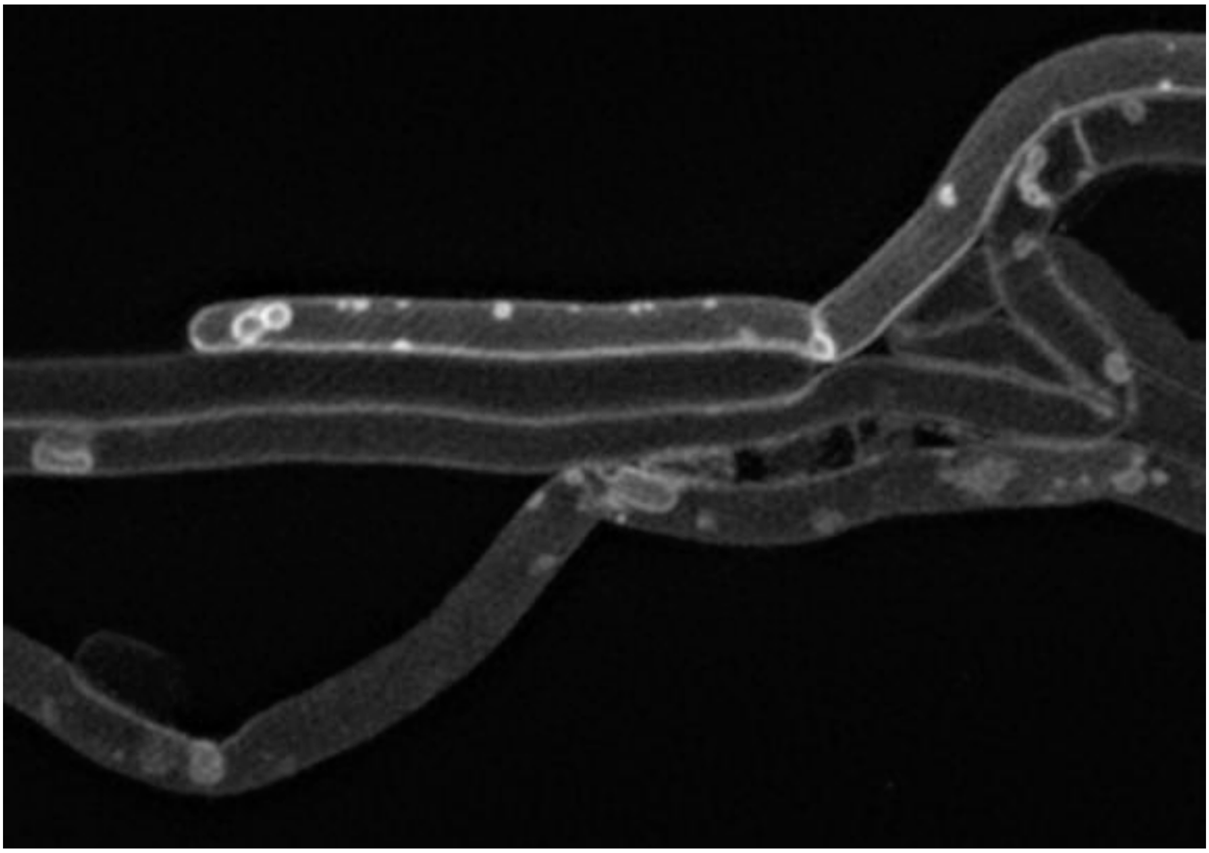

**SUPPLEMENTAL VIDEO 1:**
SIM time-lapse of BL21 DE3 containing induced pRSFDuet-1_VNp-LZ_CydAB-mNeongreen. CydAB-mNeongreen labelled inner membranes highlight dynamic movement of VNp-LZ fusion induced membrane bound cytosolic vesicles (100 msec / frame).
